# Dissection of the *Ren6* and *Ren7* powdery mildew resistance loci in *Vitis piasezkii* DVIT2027 using phased parental–progeny genomes and intraspecific locus graph reconstruction

**DOI:** 10.1101/2025.05.14.653892

**Authors:** Mélanie Massonnet, Rosa Figueroa-Balderas, Noé Cochetel, Summaira Riaz, Dániel Pap, M. Andrew Walker, Dario Cantu

## Abstract

The Chinese grape accession *Vitis piasezkii* DVIT2027 carries two loci associated with powdery mildew (PM) resistance loci, *Ren6* and *Ren7*, which differ in timing and strength of response to *Erysiphe necator*. Both loci are consistent with recognition by intracellular immune receptors. To identify the underlying nucleotide-binding leucine-rich repeat (NLR) genes, we assembled chromosome-scale diploid genomes of DVIT2027 and the susceptible *V. vinifera* F2-35, parents of a segregating F1 population. We integrated these assemblies with deep resequencing data from eight F1 sib-lines carrying different *Ren6*/*Ren7* combinations and generated trio-binned, parent-phased genomes for six progeny. This resolved both PM-resistant (PMR) and PM-susceptible (PMS) haplotypes at *Ren6* and *Ren7*. Comparative analyses revealed extensive structural variation and complete haplotype specificity among NLRs, with several candidate genes lacking allelic counterparts in PMS haplotypes. Expression profiling across PMR sib-lines identified four and two CC-NBS-LRR genes potentially associated with *Ren6* and *Ren7*, respectively. Sequence graph reconstruction of these loci across multiple *V. piasezkii* accessions revealed broad intraspecific diversity and DVIT2027-specific nodes, including within candidate NLR genes. These results provide a high-resolution view of *Ren6* and *Ren7* and support the identification of resistance gene candidates for functional validation and grapevine breeding.

## Introduction

Cultivated grapevines (*V. vinifera* spp. *vinifera*; hereafter *V. vinifera*) are generally susceptible to powdery mildew (PM), a disease caused by the obligate biotrophic ascomycete *Erysiphe necator* Schwein. (syn. *Uncinula necator*). *E. necator* infects all grapevine green tissues, including leaves, shoots, inflorescences, and berries before the onset of ripening^1^, reducing photosynthetic capacity and negatively impacting yield, fruit composition, and wine quality^2,3^. PM control relies heavily on prophylactic fungicide applications, which are costly^4^. An alternative approach is the use of resistant cultivars developed through the introgression of PM resistance loci from wild *Vitis* species and *Muscadinia rotundifolia*. To date, 15 loci associated with PM resistance have been mapped and named *Run* (Resistance to *Uncinula necator*) or *Ren* (Resistance to *E. necator*)^5–7^. Although introgression has conferred partial or complete resistance to the disease, it often introduces linkage drag from wild donor genomes^8^. The removal of unwanted wild alleles is hampered by low recombination rates in grape breeding programs and the absence of perfect markers, i.e., markers specific to the resistance (*R*) genes, due to limited knowledge of the genes underlying resistance. To date, only one *R* gene, encoding an intracellular receptor, has been characterized, at the *Run1* locus of *M. rotundifolia* G52^9^. Recent diploid genome assemblies of wild grape accessions^10^, have enabled the identification of candidate *R* genes at several loci^11^, including *Run1*.*2a*/*b* and *Run2*.*2*^12^. In this study, we leveraged recent genomic advances to identify candidate *R* genes associated with *Ren6* and *Ren7* in *V. piasezkii* DVIT2027.

Four *V. piasezkii* accessions collected during the 1980 Sino-American botanical expedition in the Shennongjia Forestry District, Hubei Province, China (DVIT2026, DVIT2027, DVIT2028, and DVIT2032) were found to be PM-resistant (PMR)^13^. Among these, only DVIT2027 has been genetically characterized. Using quantitative trait locus (QTL) mapping of the F1 population 11373, derived from a cross between the PM-susceptible (PMS) *V. vinifera* F2-35 and *V. piasezkii* DVIT2027, we previously identified two *R* loci: *Ren6* on chromosome 9 and *Ren7* on chromosome 19^13^. Both loci are heterozygous in DVIT2027 and confer resistance to *Erysiphe necator*, though with distinct response dynamics. *Ren6* provides complete resistance, while *Ren7* results in reduced colony size. In both cases, resistance relies on programmed cell death (PCD) following fungal penetration: at the appressoria of germinated spores for *Ren6*, and at the appressoria of secondary hyphae for *Ren7*. As PCD occurs post-penetration, resistance is likely triggered by intracellular recognition of *E. necator* effectors by intracellular receptors encoded within the *Ren6* and *Ren7* loci^13^.

Nucleotide-binding domain and leucine-rich repeat (NLR) proteins are intracellular receptors that activate plant defense responses upon pathogen recognition^14^. Canonical NLRs are composed of three domains: a variable N-terminal domain, a central nucleotide-binding site domain (NBS), and a C-terminal leucine-rich repeat (LRR) domain. The N-terminal domain is typically classified as coiled-coil (CC), Resistance to Powdery Mildew 8 (RPW8), or Toll/interleukin-1 receptor (TIR) type^15^. The C-terminal LRR region mediates effector recognition, either directly or indirectly, while the N-terminal domain contributes to the formation of resistosomes, large oligomeric complexes, that initiate downstream immune signaling^14,16^. NLR activation triggers calcium influx, reactive oxygen species production, and an extensive transcriptional reprogramming ^15^. It is also commonly associated with the hypersensitive response, a rapid, localized form of PCD at the site of infection^15^.

In this study, we aimed to identify NLR genes potentially responsible for PM resistance at *Ren6* and *Ren7* loci of *V. piasezkii* DVIT2027. We first generated diploid, chromosome-scale genome assemblies of *V. piasezkii* DVIT2027 and the maternal parent of the 11373 F1 population, *V. vinifera* F2-35. Since both loci are heterozygous in DVIT2027, we used genomic data from F1 progeny to distinguish *Ren6* and *Ren7* from their corresponding PMS haplotypes and to validate their structure and phasing. NLR genes within the two *R* loci and their PMS haplotypes were identified and their gene models were manually curated. To pinpoint NLRs specific to *Ren6* and *Ren7*, we compared each *R* haplotype with its PMS counterpart, focusing on structural variation affecting gene content and domain composition, and examined allelic relationships between NLRs across haplotypes. To prioritize candidate *R* genes, *i*.*e*. genes potentially involved in PM resistance, we profiled NLR gene expression in six PM-resistant 11373 sib-lines carrying different combinations of *Ren6* and *Ren7* using RNA sequencing (RNA-seq). Finally, we constructed sequence graphs for the *Ren6* and *Ren7* regions across four *V. piasezkii* accessions to assess intraspecific structural diversity and to identify graph nodes unique to DVIT2027, including those overlapping candidate NLR genes.

## Materials and Methods

### Plant material and PM inoculation

Young leaves from *V. piasezkii* DVIT2027, *V. vinifera* F2-35, and eight of their progeny ^13^: 11373-158 (*Ren6*^+^/*Ren*7^-^), 11373-195 (*Ren6*^+^/*Ren*7^-^), 11373-137 (*Ren6*^-^/*Ren*7^+^), 11373-257 (*Ren6*^-^ /*Ren*7^+^), 11373-098 (*Ren6*^+^/*Ren*7^+^), 11373-145 (*Ren6*^+^/*Ren*7^+^), 11373-130 (*Ren6*^-^/*Ren*7^-^), and 11373-156 (*Ren6*^-^/*Ren*7^-^), were collected for genome sequencing.

For gene expression profiling, PM was inoculated as described in Amrine *et al*.^17^. Three plants of each grape accession were inoculated with *E. necator* C-train, meanwhile three other plants were mock-inoculated. At 1 and 5 days post-innoculation (dpi), two leaves from each plant were pooled together to constitute a biological replicate and were immediately frozen in liquid nitrogen. Each condition was thus represented by three biological replicates.

### DNA and RNA extraction, library preparation, and sequencing

High-molecular weight genomic DNA was isolated from young leaves of *V. piasezkii* DVIT2027, *V. vinifera* F2-35, and the eight F1 progeny as in Chin *et al*.^18^. DNA quantity and purity were assessed using a Qubit™ 1X dsDNA HS Assay Kit (Thermo Fisher, MA, USA) and a Nanodrop 2000 spectrophotometer (Thermo Scientific, IL, USA), respectively. DNA integrity was evaluated with the DNA High Sensitivity kit (Life Technologies, CA, USA) and by pulsed-field gel electrophoresis. Unfortunately, the quantity and integrity of the DNA extracted from the sib-lines 11373-195 (*Ren6*^+^/*Ren*7^-^) and 11373-137 (*Ren6*^-^/*Ren*7^+^) were not sufficient for preparing long-read DNA sequencing (DNA-seq) libraries. Accordingly, DNA extracted from these two samples were only used for short-read DNA-seq. HiFi libraries of *V. piasezkii* DVIT2027 and *V. vinifera* F2-35 were prepared using the SMRTbell Express Template Prep Kit 3.0 (Pacific Biosciences, CA, USA) following the manufacturer’s protocol. Libraries were size-selected using a LightBench instrument (Yourgene Health, FL, USA) with a cut-off size range of 10 kbp, and then sequenced with a PacBio Revio platform (DNA Technology Core Facility, University of California, Davis, CA, USA). SMRTbell libraries of the 11373 sib-lines were made following the manufacturer’s protocol for the SMRTbell Express Template Prep Kit 2.0 (Pacific Biosciences, CA, USA). Size selection of the libraries was performed using a BluePippin instrument (Sage Science, Beverly, MA, USA) with a cut-off size range of 18-80 kbp. After size selection, libraries were sequenced using PacBio Sequel II system (DNA Technology Core Facility, University of California, Davis, CA, USA) (**Table S1**).

DNA extraction and short-read DNA sequencing (DNA-seq) library preparation were performed as Massonnet *et al*.^19^. DNA libraries were sequenced as 150-bp paired-end reads using an Illumina HiSeq3000 for *V. vinifera* F2-35 library, and an HiSeqX Ten system for the others (IDseq, Davis, CA, USA). Total RNA was extracted with a cetyltrimethylammonium bromide (CTAB)-based protocol as described in Cochetel *et al*.^20^. RNA purity was evaluated with a Nanodrop 2000 spectrophotometer (Thermo Scientific, IL, USA), RNA quantity with a Qubit 2.0 Fluorometer and a broad-range RNA kit (Life Technologies, CA, USA), RNA integrity through electrophoresis and with an Agilent 2100 Bioanalyzer (Agilent Technologies, CA, USA). cDNA library preparation was made as in Amrine *et al*. ^17^ using the Illumina TruSeq RNA Sample Preparation Kit v2. cDNA library of young leaves of *V. piasezkii* DVIT2027 was sequenced as 100-bp single-end reads with an Illumina HiSeq3000. Concerning the PM- and mock-inoculated leaves of the 11373 sib-lines, cDNA libraries were sequenced using an Illumina HiSeq2500 sequencer (DNA Technologies Core, University of California, Davis, CA, USA) as 50-bp single-end reads (**Table S2**). Full-length cDNA sequencing (Iso-Seq) library was prepared from young leaves of *V. piasezkii* DVIT2027 and sequenced as in Cochetel *et al*.^21^.

### Genome assembly

Long-read DNA-sequencing of the PacBio HiFi reads of *V. piasezkii* DVIT2027 and *V. vinifera* F2-35 resulted in 139 and 176X of coverage of a 500-Mbp haploid genome, respectively (**Table S1**). PacBio HiFi reads were first subsampled to 60X coverage using seqtk sample from the package seqtk v.1.2-r101-dirty (https://github.com/lh3/seqtk) and the parameter “-s100”, and then assembled using hifiasm^22^ with the option “-n 13” (**Table S3**). Contigs were scaffolded into diploid chromosome-scale pseudomolecules using HaploSync v1.0^23^ and the consensus grape synteny map (Cochetel *et al*., 2025). Telomere repeat units were searched as described in Shi *et al*.^24^, using TIDK v.0.2.63 (https://github.com/tolkit/telomeric-identifier); tidk explore with the options “--minimum 5 --maximum 12 -t 2 --log”; tidk search with the option “-s TTTAGGG”. Results were visualized with tidk plot. BUSCO v.5.4.7 with the library embryophyte_odb10^25^ were used to evaluate the gene space completeness of the genome assemblies. Concerning the six F1 individuals (11373-158, 11373-137, 11373-098, 11373-145, 11373-130, 11373-156), genome assembly of the PacBio CLR reads was performed with Canu v2.2 and the trio-binning approach with short DNA-seq reads from the two parents^26^, generating two haploid, parental-phased assemblies per F1 line. Canu was performed with the following parameters: genomeSize=500m useGrid=false maxMemory=80 maxThreads=40. For sequence polishing, PacBio CLR reads were first aligned on the genome assembly using the bioconda package pbmm2 v.1.9.0 (https://github.com/PacificBiosciences/pbmm2). Polishing was then performed with the bioconda package pbgcpp v.2.0.2-2.0.2 (https://github.com/PacificBiosciences/gcpp; downloaded August 11^th^, 2022) using the Arrow algorithm (**Table S3**).

### Genome annotation

Repetitive elements were annotated using RepeatMasker v.4.1.7-p1 (http://www.repeatmasker.org) and a grape repeat library^27^. A set of high-quality gene models was created using PASApipeline v2.5.3 (https://github.com/PASApipeline/PASApipeline), GMAP v.2024-11-20^28^, and pblat v.2.5.1^29^. For the gene annotation of *V. piasezkii* DVIT2027, we used as transcriptomic evidence the PN40024 V5.1 predicted protein-coding sequences (https://grapedia.org/t2t_annotation/) and the high-quality isoforms of *V. piasezkii* DVIT2027. For *V. vinifera* F2-35, transcript evidences consisted in the PN40024 V5.1 predicted protein-coding sequences and the high-quality isoforms of Cabernet Sauvignon^27^. Transcripts were extracted from the Iso-Seq reads of *V. piasezkii* DVIT2027 young leaves like Cochetel *et al*.^21^. The set of high-quality gene models was used as input for the *ab initio* predictors Augustus v.3.5.0^30^ and GeneMark v. 3.68_lic^31^. EvidenceModeler v.2.1.0^32^ was then used to generate consensus models from the gene models produced by the *ab initio* predictions and PASA. Gene models encoding protein sequences with a length inferior to 50 amino acids, in-frame stop codons, no start methionine, no stop codon, were removed.

### Functional annotation

Predicted proteins of *V. piasezkii* DVIT2027 and *V. vinifera* F2-35 genomes were aligned onto the proteins of *Arabidopsis thaliana* (Araport11_genes.201606.pep.fasta; https://www.arabidopsis.org/download/index.jsp) using BLASTP v.2.12.0+^33^. Alignments with an identity greater than 30%, and both reference:query and query:reference coverages between 75% and 125%, were retained. Among the remaining alignments, the alignment with the highest product of identity, query coverage, and reference coverage, was selected for each grape protein to determine its homolog in *A. thaliana*. Same methods were used to identify homologous proteins in the grape *V. vinifera* PN40024 genome V1 and V5.1 annotations^24,34^.

Protein domains were detected using hmmsearch from HMMER v.3.3.1 (http://hmmer.org/) and the Pfam-A Hidden Markov Models (HMM) database^35^ (downloaded on March 12^th^, 2025). Protein domains with an independent E-value < 1.0, and at least 50% of the HMM, were selected. CC domains were identified using COILS^36^. Functional annotation of the predicted proteins of *V. piasezkii* DVIT2027 and *V. vinifera* F2-35 genomes can be retrieved in **Tables S4-S5**. LRR motifs among the candidate NLRs were predicted using LRRpredictor v1.0^37^. The CC, NBS, and LRR domains identified among the NLRs composing *Ren6* and *Ren7* were used to manually annotate them in the genome of *V. piasezkii* DVIT2027.

### NLR gene annotation

Gene loci potentially encoding NLRs were detected among *V. piasezkii* DVIT2027 and *V. vinifera* F2-35 genomes with NLR-Annotator using default parameters^38^. To assess the intron-exon structure of the potential NLR genes within the *R* loci and the PMS alternative haplotypes, RNA-seq reads from *V. piasezkii* DVIT2027, *V. vinifera* F2-35, and their eight progeny lines were aligned on the combined genomes of *V. piasezkii* DVIT2027, *V. vinifera* F2-35, and *E. necator* C-strain^39^, using HISAT2 v.2.1.0^40^ and the parameters “--end-to-end --sensitive -k 50”. RNA-seq reads of *V. vinifera* F2-35 were retrieved from the NCBI BioProject PRJNA897013^41^. Alignments were visualized in Integrative Genomics Viewer (IGV) v.2.4.14^42^. Manual refinement of the NLR gene models was performed when the alignments of the RNA-seq reads indicated a different intron-exon structure than the *ab initio* structural annotation.

### *Ren6* and *Ren7* region localization

The loci *Ren6* and *Ren7* and their alternative haplotypes were located by *in silico* amplification of the *Ren6*- and *Ren7*-associated markers (**Table S6**) described in Pap *et al*.^13^ on the genome assemblies using dipr (https://github.com/douglasgscofield/dispr).

### Haplotype coverage analysis

To distinguish *Ren6* and *Ren7* haplotypes from their PMS alternative haplotypes and confirm their phasing in the genomes of *V. piasezkii* DVIT2027 and *V. vinifera* F2-35, we used the short DNA-seq reads from the eight F1 lines, *V. piasezkii* DVIT2027 and *V. vinifera* F2-35. Adapter sequences and low-quality reads were removed using Trimmomatic v.0.36^43^ with the following parameters: “LEADING:3 TRAILING:3 SLIDINGWINDOW:10:20 MINLEN:36 CROP:150”. High-quality reads were then aligned onto the combined genomes of *V. piasezkii* DVIT2027 and *V. vinifera* F2-35 using BWA v0.7.17^44^ with default parameters. Reads aligning perfectly (i.e., with no mismatches) to the reference genome were selected with bamtools filter v.2.5.1^45^ and the option “NM:0”, ensuring that only unambiguous alignments were retained. This strict criterion was applied to prevent reads carrying parental SNPs from mapping to both haplotypes, allowing accurate discrimination of parental haplotype blocks and reliable estimation of haplotype-specific coverage. Base coverage was performed using genomecov from BEDTools v2.29.1^46^. Coverage in repetitive elements was removed using BEDTools intersect. Median coverage per window of 10-kbp was computed by BEDTools map. For each grape accession, the median coverage per 10-kbp window was normalized by dividing by the sequencing coverage of the accession (**Table S2**). Phasing and structure of the *Ren6* and *Ren7* haplotypes and their alternative haplotypes in the genome of *V. piasezkii* DVIT2027 were also confirmed using the alignment of the PacBio CLR reads of six of the eight sib-lines on the four haplotypes. Reads were first subsampled to 40X coverage using seqtk sample from the package seqtk v.1.2-r101-dirty (https://github.com/lh3/seqtk) and the parameter “-s100”, and then aligned onto the combined genomes of *V. piasezkii* DVIT2027 and *V. vinifera* F2-35 with Minimap2 v.2.24-r1122^47^. Alignments were visualized using IGV v.2.4.14^42^.

### Sequence comparison

Pairwise sequence alignments of whole haplotypes were done with NUCmer from MUMmer v.4.0.0^48^ with the “--mum” option. For the alignment of 11373 sib-lines’ genomes onto *Ren6, Ren7*, and their alternative haplotypes, each parental genome was aligned onto the parental haplotypes. Alignments longer than 5 kbp were retained using the parameters “-l 5000”. Regarding the comparison between PMR and PMS haplotypes, alignments with at least 90% identity are represented in **Figure 2**. Structural variants (SVs; > 50 bp) were called using show-diff from MUMmer v.4.0.0 with defaults parameters. SNPs and INDELs (< 50 bp) were detected using show-snps from MUMmer v.4.0.0. SV coordinates were intersected with the CDS annotation of the NLRs composing *Ren6* and *Ren7* using BEDTools intersect v2.29.1^46^ to determine the impact of the SVs on the NLR genes. SVs affecting the NLR gene content of *Ren6* and *Ren7* were manually inspected using the alignment results. The potential effect of the SNPs, detected between *Ren6* and *Ren7* and their corresponding PMS alternative haplotype in DVIT2027, on the amino acid content was predicted using SnpEff v.4.3t^49^.

To assess whether NLR genes of *Ren6* and *Ren7* loci were present in *V. piasezkii* 0957, BL-1 and BS-40^50,51^, CDS of the NLR genes composing the two *R* loci were first aligned onto the four haploid genomes using GMAP v.2020-06-01^28^ (**Tables S7-S9**). Resulting alignments were then used to generate proteins by gffread from Cufflinks v.2.2.1^52^. Sequences of the latter ones were compared to the protein sequences of the NLRs composing *Ren6* and *Ren7* using BLAST v.2.2.28+^33^.

### Phylogenetic analysis

Predicted NLR protein sequences from *Ren6* and *Ren7*, and their PMS alternative haplotypes in *V. piasezkii* DVIT2027 and *V. vinifera* F2-35 were aligned using MUSCLE^53^ in MEGAX^54^. Phylogenetic analyses were done with MEGAX^54^ using the Neighbor-Joining method^55^ and 1,000 bootstrap replicates. Evolutionary distances were computed with the Poisson correction method^56^, eliminating the positions with less than 5% site coverage.

### Gene expression analysis

High-quality reads were obtained using Trimmomatic v.0.36^43^ with the following parameters: “LEADING:3 TRAILING:3 SLIDINGWINDOW:10:20 MINLEN:36” (**Table S10**). Transcript abundance was evaluated with Salmon v.1.5.1^57^. First, the transcriptome index file was built using the combined protein-coding sequences of *V. piasezkii* DVIT2027 and *V. vinifera* F2-35, and *E. necator* C-strain^39^, their genomes as decoys, and the options: “-k 13 --keepDuplicates”. Second, transcript abundance was quantified with the parameters “--gcBias –seqBias --validateMappings”. Read counts from *E. necator* C-strain were removed, and gene expression in transcript per million (TPM) from the grape CDS were calculated. Differential gene expression analysis was performed using DESeq2 v.1.42.1^58^ after importing the quantification files with the R package tximport v.1.30.0^59^.

### Sequence graph construction

Sequence graphs of *Ren6* and *Ren7* and their respective six alternative haplotypes were built using the nf-core/pangenome pipeline^60^ with NextFlow v.25.04.6^61^ and the parameters “-r 1.1.2 -profile singularity”.

## Results

### Haplotype-resolved genome assemblies of *V. piasezkii* DVIT2027 and *V. vinifera* F2-35

To identify *R* genes potentially involved in PM resistance of *V. piasezkii* DVIT2027, we sequenced the genomes of the paternal (*V. piasezkii* DVIT2027) and maternal (*V. vinifera* F2-35) parents of the 11373 F1 population^13^, using long-read DNA sequencing (PacBio HiFi reads; **Table S1**). Draft haplotype-resolved genome assemblies comprised 120 contigs for *V. piasezkii* DVIT2027 and 238 contigs for *V. vinifera* F2-35, totaling 1,018.8 Mbp and 975.7 Mbp, respectively (**Table S3**). Pseudomolecule reconstruction yielded two sets of 19 chromosomes: 504.6 Mbp (Hap1) and 507.1 Mbp (Hap2) for DVIT2027, and 481.3 Mbp (Hap1) and 481.5 Mbp (Hap2) for F2-35 (**Table 1**). The difference of length between the two genome assemblies (38.1 Mbp) was mostly due to longer chromosomes 12,13, 16, 17, and 18 in DVIT2027 compared to F2-35 (**Table S11**). This size difference was made of repetitive elements (49.2%), but also of protein-coding gene loci (12.3%).

**Table 1:** Statistics of the genome assemblies of *V. piasezkii* DVIT2027 and *V. vinifera* F2-35.

|  | *V. piasezkii* DVIT2027 |  |  | *V. vinifera* F2-35 |  |  |
| --- | --- | --- | --- | --- | --- | --- |
|  | Hap 1 | Hap 2 | Unplaced | Hap 1 | Hap 2 | Unplaced |
| Assembly length (Mbp) | 504.6 | 507.1 | 7.6 | 481.3 | 481.5 | 18.4 |
| Number of sequences | 19 | 19 | 55 | 19 | 19 | 93 |
| Average length (Mbp) | 26.6 | 26.7 | 0.1 | 25.3 | 25.3 | 0.2 |
| Telomeres | 31/38 | 33/38 | 0 | 34/38 | 33/38 | 0 |
| Number of gene loci | 34,926 | 31,353 | 652 | 29,820 | 29,399 | 394 |
| Gene content (Mbp) | 149.6 | 134.3 | 1.7 | 141.9 | 138.1 | 0.9 |
| Repeats (Mbp) | 284.1 | 285.0 | 5.5 | 270.0 | 270.0 | 15.8 |
| Repeats (%) | 56.3 | 56.3 | 71.9 | 56.1 | 56.1 | 85.9 |
| BUSCO (%) | 99.3 | 99.5 | 0.6 | 99.4 | 99.1 | 0.2 |

Unanchored contigs accounted for only 0.7% (DVIT2027) and 1.9% (F2-35) of the total genome size and were predominantly composed of repetitive sequences (**Table 1**). BUSCO analysis showed high completeness, with 99.3 ± 0.2% of expected single-copy orthologs identified in each haplotype. On average, 32.8 ± 1.3 telomeres of the 38 possible telomeres were detected per haplotype. Together, these results demonstrate that we generated highly contiguous, haplotype-resolved genome assemblies for both parents of the 11373 population.

### Identification and haplotype phasing of *Ren6* and *Ren7* loci of *V. piasezkii* DVIT2027

The boundaries of the *Ren6* and *Ren7* haplotypes, along with their PMS alternative haplotypes, were defined by *in silico* amplification of the *Ren6*-associated markers PN9-057 and VMC4h4.1, and *Ren7*-associated markers VMC9a2.1 and VVIu09^13^ on the diploid genomes of DVIT2027 and F2-35. Because *Ren6* and *Ren7* are heterozygous in DVIT2027, we distinguished the PMR haplotypes from their PMS alternatives, and validated haplotype phasing by analyzing normalized median base coverage (per 10 kbp) across both parental genomes using deep sequencing data of eight sib-lines of population 11373^13^, including: two lines carrying *Ren6* only (*Ren6*^+^/*Ren7*^-^: 11373-158 and 11373-195), two with *Ren7* only (*Ren6*^-^/*Ren7*^+^: 11373-137 and 11373-257), two carrying both loci (*Ren6*^+^/*Ren7*^+^: 11373-098 and 11373-145), and two lacking both (*Ren6*^-^/*Ren7*^-^: 11373-130 and 11373-156). In *Ren6*^+^ accessions, short-read coverage was restricted to haplotype 1 of chromosome 9, whereas *Ren6*^-^ accessions showed coverage only on haplotype 2 (**Figures S1-S3**). Similarly, for *Ren7*^+^ lines, coverage was confined on haplotype 1 of chromosome 19, while *Ren7*^-^ lines mapped to haplotype 2 (**Figures S2-S4**). These results indicate that *Ren6* and *Ren7* are located on haplotype 1 of chromosome 9 and 19, respectively, and their PMS counterparts, *i*.*e*. PMS alternative haplotypes, reside on haplotype 2. They also indicate no haplotype switching occurred during the assembly of DVIT2027. In F2-35, coverage profiles were consistent with the inheritance of PMS haplotypes in the F1 sib-lines, with no evidence of recombination or haplotype switching (**Figures S5-S8**).

To further validate the phasing, structure, and sequence of the *Ren6* and *Ren7* loci and their PMS alternative haplotypes, we sequenced the genomes of six of the eight sib-lines using PacBio CLR reads. Alignment of the long DNA-seq reads onto the genomes of DVIT2027 and F2-35 confirmed correct haplotype phasing. Consistent and even read coverage across these regions indicated no misassemblies that could compromise sequence structure (**Figures S9-10**). We then assembled the six sib-line genomes using a trio-binning approach^26^, generating two haploid, parental-phased assemblies per line: one inherited from DVIT2027 and the other from F2-35 (**Table S3**). Alignment of these assemblies to the *Ren6, Ren7*, and PMS haplotypes showed complete coverage of the corresponding loci in all six sib-lines (**Figure 1**). The only exception was the *Ren6*-PMS haplotype of DVIT2027, which was not fully spanned by a single contig in one sib-line. Overall, the alignments showed high sequence similarity, with an average identity of 99.96 ± 0.04% (**Figure 1**).

**Figure 1:**
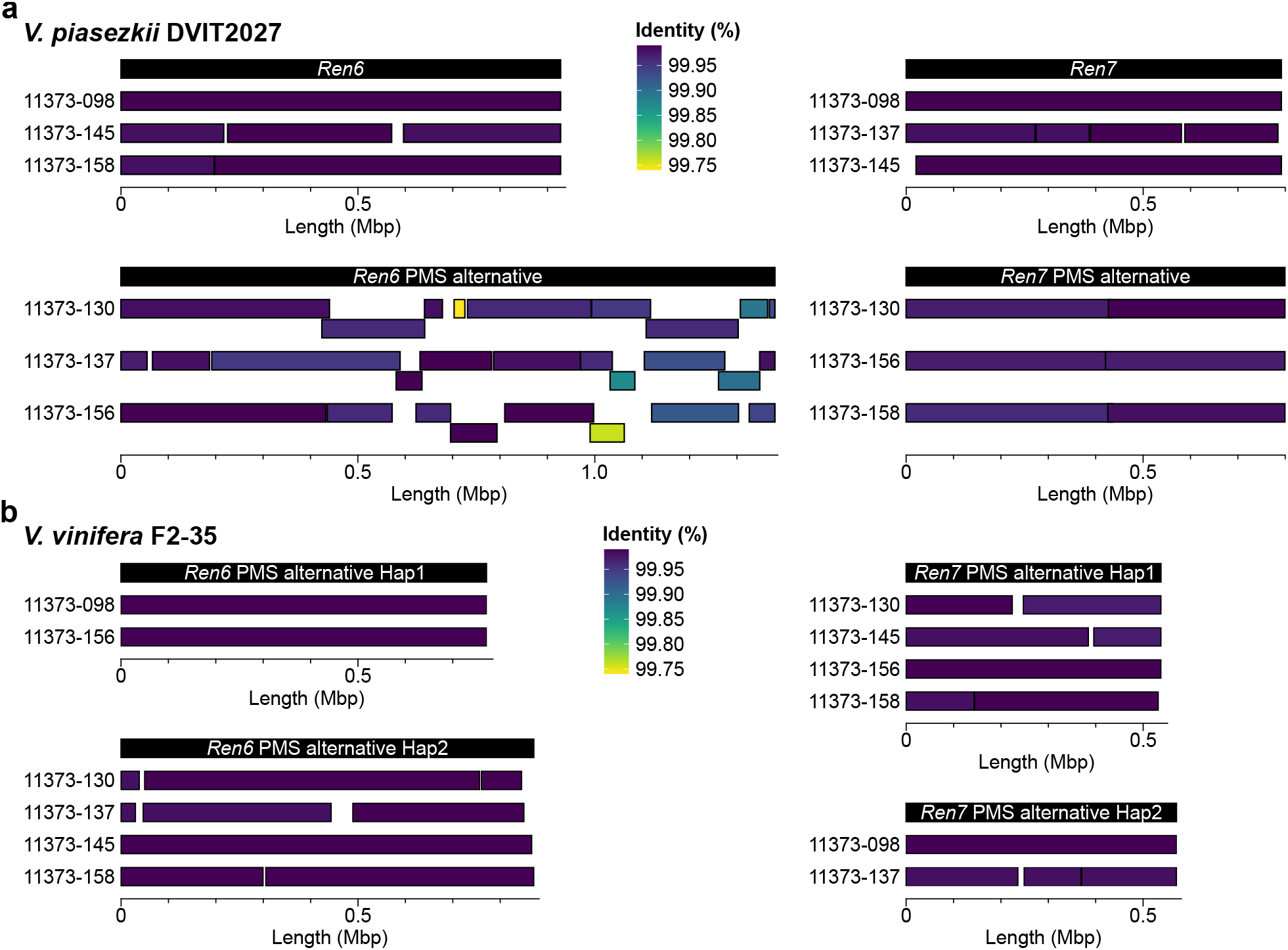
Validation of the haplotype phasing and structure of *Ren6* and *Ren*7 loci and their PM-susceptible alternative haplotypes using progeny genomes. Alignment of the genome assemblies of six 11373 sib-lines onto *Ren6, Ren7*, and their PM-susceptible (PMS) alternatives from *V. piasezkii* DVIT2027 (**a**) and *V. vinifera* F2-35 (**b**). Each square represents a contig sequence.

Together, these results validate the phasing and structure of the haplotypes in both parents, *V. piasezkii* DVIT2027 and *V. vinifera* F2-35. In the DVIT2027 genome, the *Ren6* and *Ren7* loci measured 927.5 and 791.6 kbp, respectively, while their PMS alternative haplotypes were 1,380.1 and 771.1 kbp (**Table 2**). In F2-35, the PMS alternative haplotypes corresponding to *Ren6* and *Ren7* were shorter than those of DVIT2027, measuring 771.1 and 870.7 kbp for *Ren6*-PMS haplotypes, and 537.4 and 570.3 kbp for *Ren7*-PMS haplotypes (**Figure 1**).

**Table 2:** Characteristics of the *Ren6* and *Ren7* loci and their alternative PM-susceptible (PMS) haplotypes in *V. piasezkii* DVIT2027 and *V. vinifera* F2-35.

| Grape accession | *V. piasezkii* DVIT2027 |  |  |  | *V. vinifera* F2-35 |  |  |  |
| --- | --- | --- | --- | --- | --- | --- | --- | --- |
| Locus | *Ren6* | *Ren6*-PMS alternative | *Ren7* | *Ren7*-PMS alternative | *Ren6*-PMS alternative Hap1 | *Ren6*-PMS alternative Hap2 | *Ren7*-PMS alternative Hap1 | *Ren7*-PMS alternative Hap2 |
| Chromosome | Chr09 Hap1 | Chr09 Hap2 | Chr19 Hap1 | Chr19 Hap2 | Chr09 Hap1 | Chr09 Hap2 | Chr19 Hap1 | Chr19 Hap2 |
| Start-Stop (bp) | 8,626,384-9,553,923 | 9,061,939-10,442,008 | 883,779-1,675,391 | 951,279-1,750,630 | 8,690,011-9,461,149 | 8,080,301-8,951,049 | 904,827-1,442,232 | 843,589-1,413,925 |
| Length (bp) | 927,540 | 1,380,070 | 791,613 | 799,352 | 771,139 | 870,749 | 537,406 | 570,337 |
| Protein-coding gene loci | 54 | 80 | 81 | 77 | 42 | 50 | 49 | 56 |
| Total NLR genes | 19 | 35 | 13 | 8 | 12 | 12 | 2 | 3 |
| CC-NBS-LRR | 15 | 25 | 10 | 6 | 10 | 8 | 2 | 2 |
| CC-NBS | 2 | 5 | 0 | 1 | 1 | 2 | 0 | 0 |
| NBS-LRR | 2 | 4 | 3 | 1 | 1 | 2 | 0 | 1 |
| NBS | 0 | 1 | 0 | 0 | 0 | 0 | 0 | 0 |

### Identification of the NLR genes composing *Ren6, Ren7*, and their PMS alternative haplotypes

Gene annotation, combined with NLR gene prediction and manual refinement of gene models, revealed differences in gene content, particularly NLR genes, between the *R* loci and their PMS counterparts (**Table 2**). Nineteen and thirteen NLR genes were annotated within the *Ren6* and *Ren7* loci, respectively. In *Ren6*, two NLR genes were located between markers PN9-063 and PN9-066.1 (202.6 kbp), while the remaining NLR genes clustered between PN9-067 and VMC4h4.1 (664.3 kbp) (**Figure 2a**). *Ren6* was mapped between PN9-063 and PN9-067.2 based on QTL analysis^13^; however, PN9-067.2 and PN9-068 amplified at multiple locations, complicating locus delimitation and reducing the resolution for identifying candidate *R* genes (**Table S12**). Notably, the *Ren6*-PMS haplotype of DVIT2027 contained more NLR genes than the PMR haplotype (35 *vs*. 20), with the NLR-containing region spanning 1,209.3 kbp compared to 747.9 kbp in the PMR haplotype, accounting for the observed length difference. In contrast, *Ren7* contained more NLR genes in the PMR haplotype than in the PMS counterpart (13 *vs*. 8). These NLRs were clustered within 266.2 kbp and 183.9 kbp regions, respectively, located between markers VMC9a2.1 and VMC5h11 (locus spans of 606.5 kbp and 621.4 kbp, respectively) (**Figure 2b**), consistent with the previously defined QTL boundaries^13^. In both *Ren6* and *Ren7*, most NLR genes encoded CC-NBS-LRR proteins (**Table 2**; **Table S13**). In F2-35, twelve NLR genes were annotated in each *Ren6*-PMS haplotype, while only two and three NLR genes were identified in the *Ren7*-PMS haplotypes.

**Figure 2:**
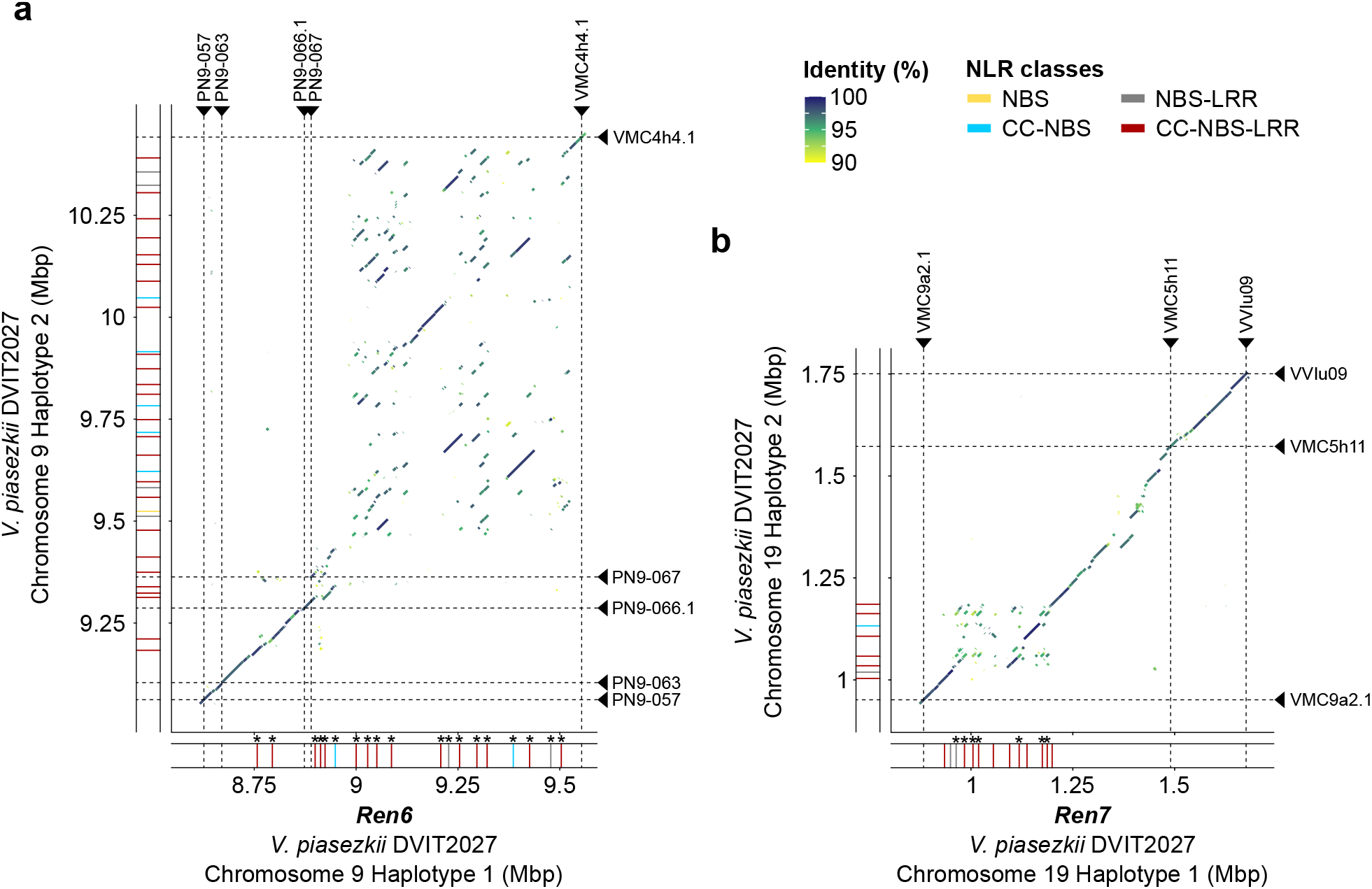
Effect of the structural variations between *Ren6* and *Ren7* loci and their PM-susceptible alternative haplotypes on the NLR gene content. Alignment of the PM-susceptible alternative haplotypes of *Ren6* (**a**) and *Ren7* (**b**) from *V. piasezkii* DVIT2027 onto their PM-resistant counterparts. Chromosomal positions of the NLR genes composing the four haplotypes are represented by colored rectangles, which color indicates the NLR class. NLR genes in *Ren6* and *Ren7* which protein-coding sequence is impacted by a structural variation are indicated by an asterisk. Panels **a** and **b** share the same legend.

### Comparison of *Ren6* and *Ren7* loci with their PM-susceptible alternative haplotypes

To identify NLRs potentially associated with *Ren6* and *Ren7* PM resistance, we compared the PMR haplotypes with their respective PMS counterparts. First, we identified structural variations (SVs; > 50bp). In *Ren6*, SVs between markers PN9-063 and PN9-066.1 were relatively small (1.0 ± 1.6 kbp) with the region spanning 202.6 kbp in the PMR haplotype and 183 kbp in the PMS haplotype (**Figure 2a**). In contrast, numerous large SVs (>10 kbp), particularly duplications, were detected between PN9-067 and VMC4h4.1. This region spanned 664.3 kbp in the PMR haplotype and 1,078.9 kbp in the PMS haplotype, largely accounting for the observed length difference. Manual inspection of the SVs affecting the coding sequences (CDS) of the NLR genes of *Ren6* revealed that all of them were entirely and/or partially duplicated in DVIT2027’s PMS alternative haplotype. By annotating the NLR domains in the genome, we found that these duplication events altered the domain composition of 13 NLRs (**Figure 2a**; **Table S14**).

For *Ren7*, a large complex SV spanning 133.6 kbp (from 955.0 to 1,088.6 kbp) was detected on haplotype 1 of chromosome 19 in DVIT2027, relative to haplotype 2 (**Figure 2b**). This SV disrupted the CDS of 4 NLR genes (**Table S15**), mainly accounting for the difference in NLR gene number between the two haplotypes. Duplication events were identified for 7 NLR genes affecting the domain composition of 6 of them (**Table S15**). Comparable SVs were also identified when comparing the PMS haplotypes of F2-35 with the *Ren6* and *Ren7* loci (**Figure S12**).

In addition to SVs, we identified small polymorphisms, including 46,178 and 20,669 SNPs and 6,629 and 3,928 INDELs (< 50 bp) in *Ren6* and *Ren7*, respectively (**Tables S16-S17**). Comparison with the CDS of the NLR genes showed that all the NLRs in *Ren6* and *Ren7* were affected by small polymorphisms. Prediction of the effect of the SNPs showed that every NLRs from *Ren6* and *Ren7*, except g332400 from *Ren7* in which only an INDEL was detected, were affected by at least one non-synonymous SNP (from 7 to 1,079 in *Ren6*, and from 1 to 376 in *Ren7*) (**Tables S18-S19**). Among the non-synonymous SNPs, 77 and 33 ones were found to lead to a premature stop codon in 14 and 7 NLRs of *Ren6* and *Ren7*, respectively. These results suggest that the two loci are highly polymorphic.

To further assess the impact of SVs and small polymorphisms on NLR gene content, we analyzed allelic relationships by constructing phylogenetic trees for the NLR genes at each *R* locus (**Figure 3**). In DVIT2027, 2 of the 19 NLR genes (10.5%) in *Ren6* (g158080 and g158100) and 6 of the 13 NLR genes (46.1%) in *Ren7* (g332310, g332320, g332340, g332350, g332390, g332430) were hemizygous, *i*.*e*. they lacked allelic counterparts in their respective PMS haplotypes (**Tables S14-S15**). While some of these genes have allelic counterparts in F2-35, their absence in the PMS haplotypes of DVIT2027 suggests that they may contribute to *Ren6*- and *Ren7*-mediated PM resistance. Noteworthily, no allele pairs shared identical protein sequences, indicating that all NLR genes within *Ren6* and *Ren7* are unique to the PMR haplotypes and distinct from their PMS counterparts in DVIT2027 and F2-35.

**Figure 3:**
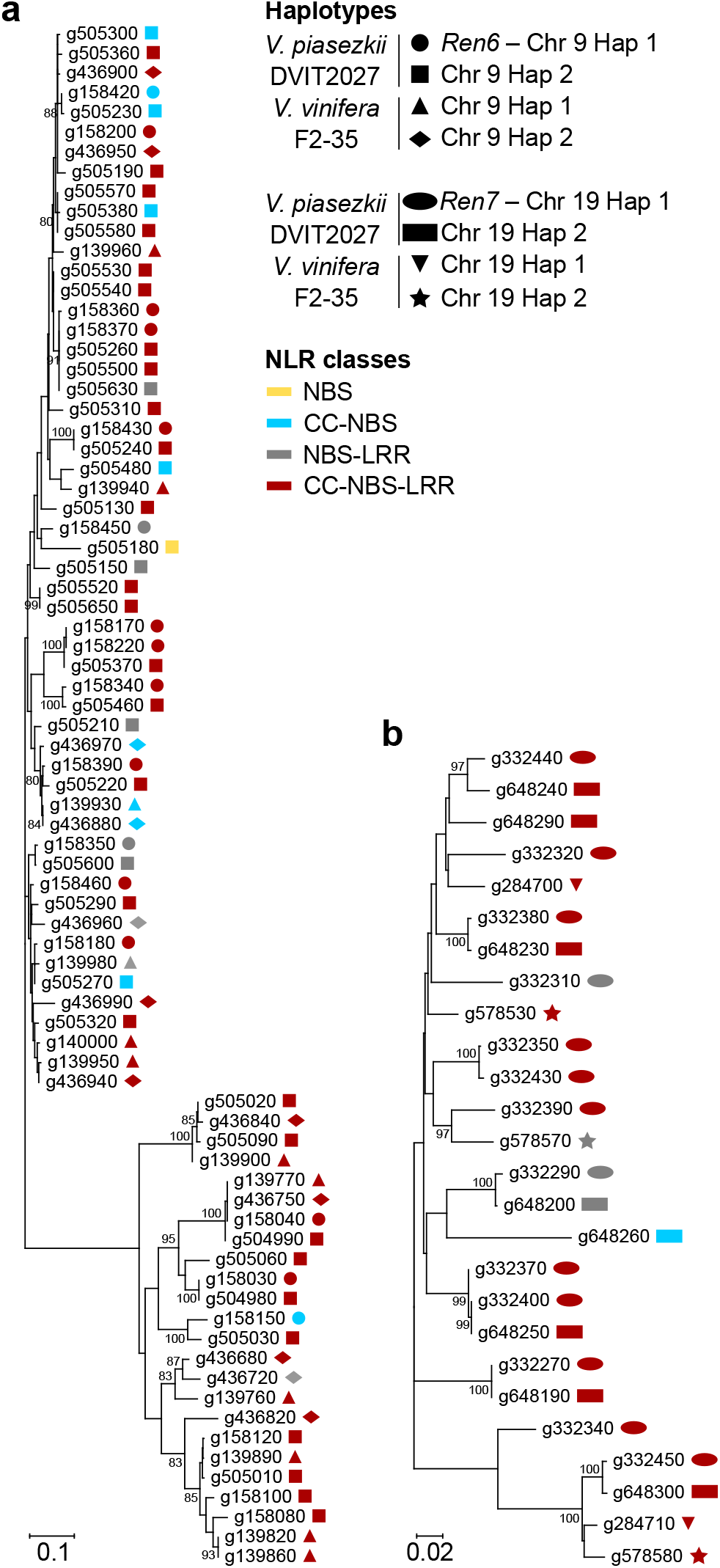
Identification of allele relationship between NLRs composing *Ren6* and *Ren7*, and their PM-susceptible alternative haplotypes, based on phylogeny. Neighbor-joining clustering of the predicted NLR sequences composing *Ren6* (**a**) and *Ren7* (**b**), and their PM-susceptible alternative haplotypes. Scale bar is in the unit of the number of substitutions per site. Bootstrap values greater than 80 are indicated. Panels **a** and **b** share the same legend.

Together, these results demonstrate that SVs and small polymorphisms affect both the number and sequence of NLR genes at *Ren6* and *Ren7*, supporting the candidacy of all NLRs in these loci as potential contributors to PM resistance.

### Identification of expressed NLR genes within *Ren6* and *Ren7*

To narrow down the NLR genes potentially responsible for *Ren6*- and *Ren7*-mediated PM resistance, we analyzed their expression in leaves of six 11373 sib-lines carrying *Ren6, Ren7*, or both loci. Leaves were sampled one and five days after inoculation with *E. necator* or a mock solution. Within *Ren6*, four CC-NBS-LRR genes were consistently expressed (>0 TPM) in all *Ren6*^+^ accessions, both constitutively and in response to *E. necator*. One gene (g158030) was located between markers PN9-063 and PN9-066.1, while the other three (g158080; g158100; g158430) were located between PN9-067 and VMC4h4.1 (**Figure 4a**). Interestingly, two of these genes (g158080 and g158100) lack allelic counterparts in the PMS haplotype of DVIT2027. For *Ren7*, four CC-NBS-LRR genes were consistently expressed (>0 TPM) in all the *Ren7*^+^ accessions. Two of them, g332320 and g332350, were expressed at a very low level (0.3 ± 0.1 and 0.2 ± 0.1 TPM, respectively), both lacked allelic counterpart in the PMS haplotype. In contrast, the two other CC-NBS-LRR genes, g332380 and g332390, showed higher expression levels (2.7 ± 0.1 and ± 1.6 TPM, respectively), with g332390 also missing an allelic counterpart in the PMS haplotype.

**Figure 4:**
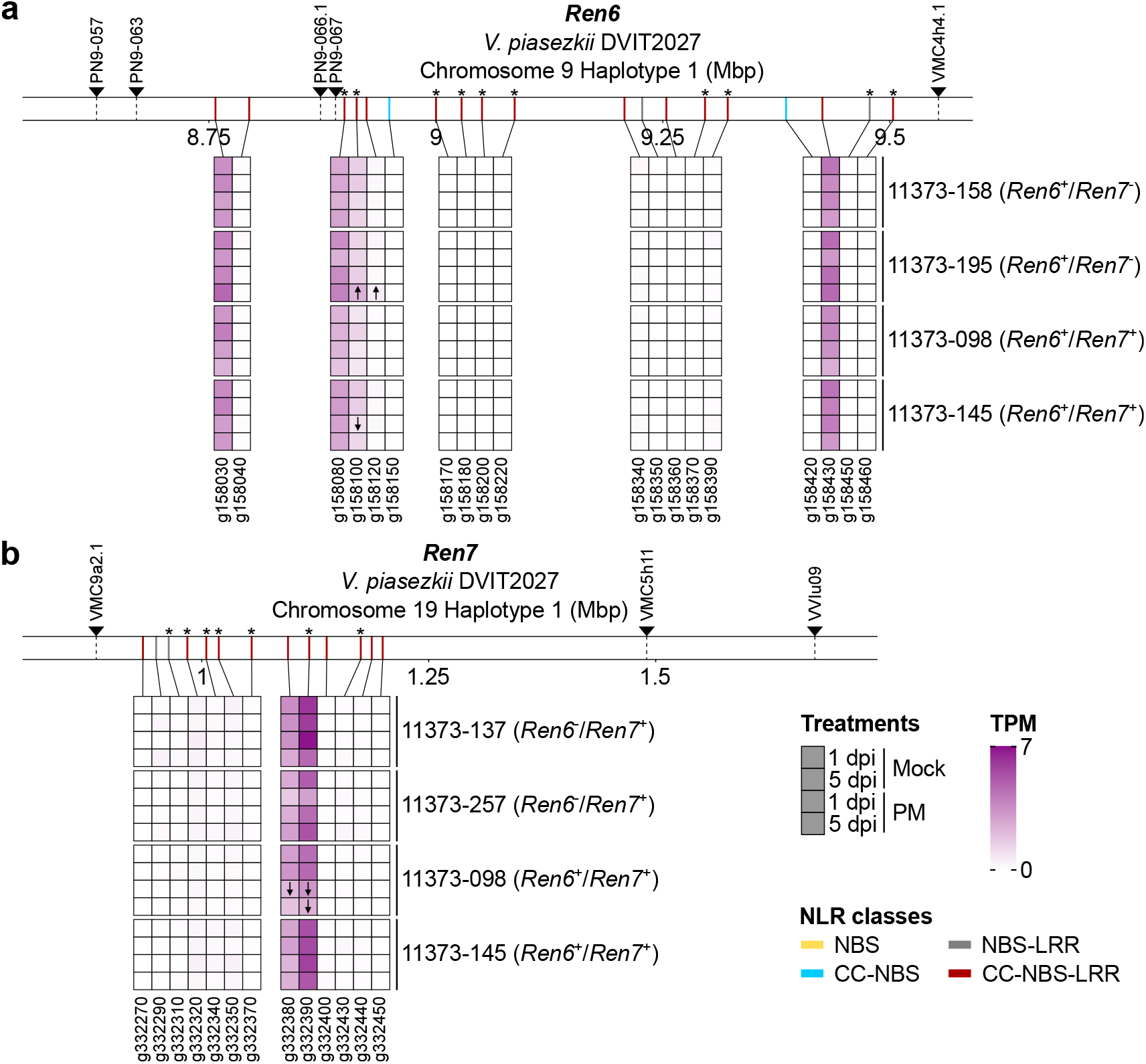
Identification of candidate NLRs responsible for PM resistance within *Ren6* and *Ren7* based on constitutive gene expression. Transcript abundance of the NLR genes belonging to *Ren6* (**a**) and *Ren7* (**b**) loci. Gene expression was measured in 11373 sib-lines possessing either *Ren6, Ren7*, or both *R* loci, at one and five days post-inoculation (dpi) with a mock solution (Mock) or *E. necator* conidia (PM). For each condition, transcript abundance is represented as the mean of transcript per million (TPM) of three bioreplicates (n = 3). NLR genes more highly and lowly expressed in response to PM are indicated by ↑ and ↓, respectively. Chromosomal positions of the NLR genes composing the four haplotypes are represented by colored rectangles, with the color indicating the NLR class. NLR genes which protein-coding sequence is impacted by at least one structural variation relative to the alternative PMS haplotype of *V. piasezkii* DVIT2027 are indicated by an asterisk. Panels **a** and **b** share the same legend.

To determine the transcriptional modulation of NLRs in *Ren6* and *Ren7* in response to PM, we performed differential gene expression analysis. No NLR gene was differentially expressed in response to PM (*P* value < 0.05) in all *Ren6*^+^ or *Ren7*^+^ accessions (**Figure 4**). However, within *Ren6*, expression of two CC-NBS-LRR genes (g158100 and g158120) were significantly induced (*P* value < 0.05) in response to PM in 11373-195 (*Ren6*^+^/*Ren7*^-^) and 11373-145 (*Ren6*^+^/*Ren7*^+^) (**Figure 4a**). In *Ren7*, two CC-NBS-LRR genes (g332380 and g332390) were significantly downregulated (*P* value < 0.05) in 11373-098 (*Ren6*^+^/*Ren7*^+^) following *E. necator* inoculation (**Figure 4b**).

Altogether, our results identify four (g158030; g158080; g158100; g158430) and two (g332380 and g332390) candidate CC-NBS-LRR genes potentially associated with *Ren6* and *Ren7* resistance, respectively, based on their consistent expression in PMR sib-lines, and their absence or sequence divergence in the corresponding PMS haplotype (**Figure 5**). Notably, g158080 and g158100 (*Ren6*) and g332390 (*Ren7*) lack allelic counterparts in the PMS haplotype of DVIT2027. The proteins encoded by g158030 and g158430 (*Ren6*) differ from their PMS alleles by two and one amino acid substitutions, respectively, including one in the CC domain of g158030 (**Figure 5a**). For g332380, the predicted protein differs from its PMS allele by two amino acid substitutions, including one in the NB-ARC domain, and two deletions (5 and 91 amino acids) at the C-terminus (**Figure 5b**).

**Figure 5:**
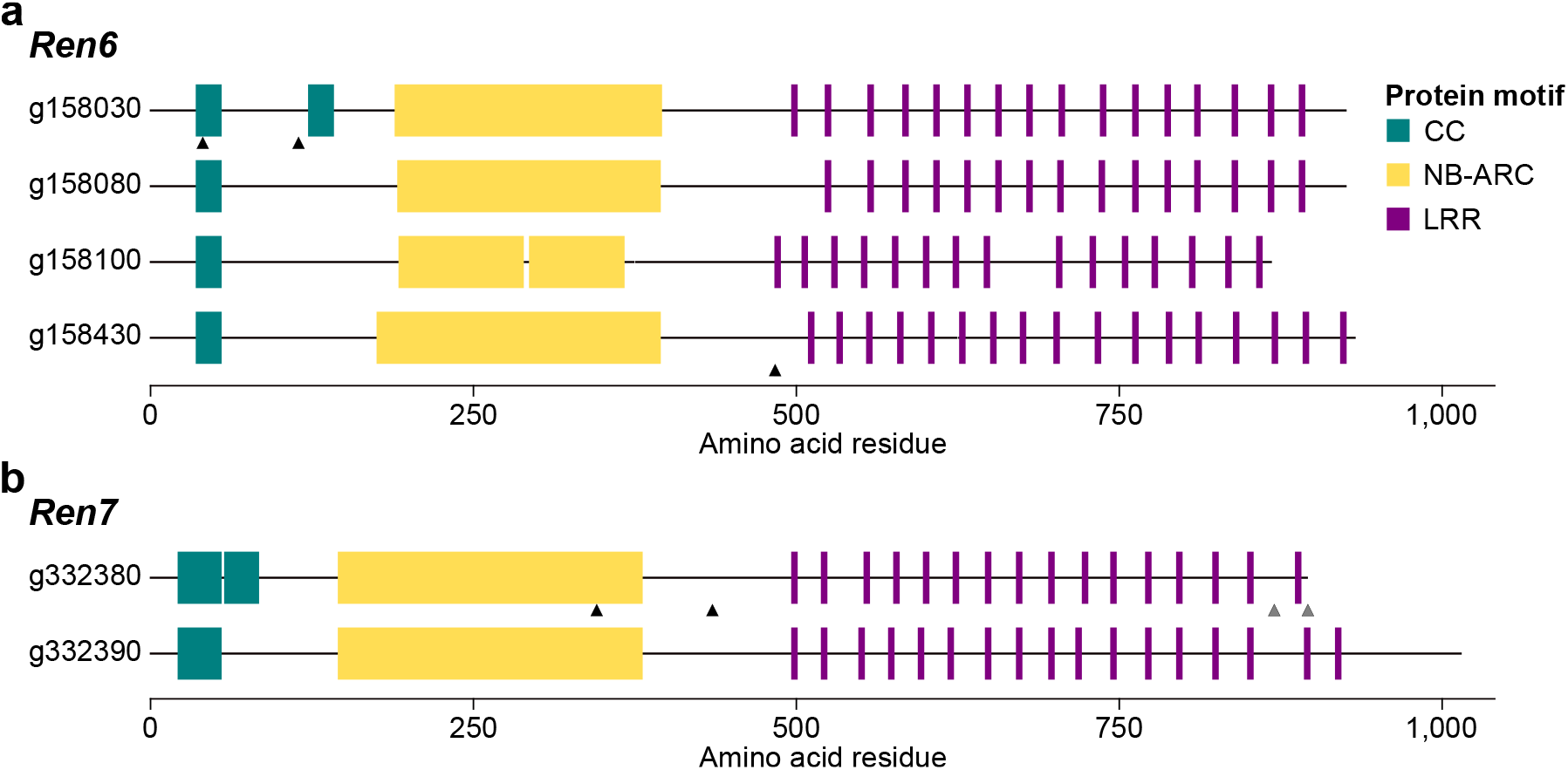
Protein motif composition of the candidate CC-NBS-LRRs associated with the PM resistance of *Ren6* (a) and *Ren7* (b) of *V. piasezkii* DVIT2027. Black and gray triangles indicate the position of the amino changes and deletions compared to the allelic counterpart located in the PM-susceptible haplotype of *V. piasezkii* DVIT2027, respectively. Panels **a** and **b** share the same legend.

### Comparison of *Ren6* and *Ren7* haplotypes across multiple *V. piasezkii* accessions

We compared the *Ren6* and *Ren7* in *V. piasezkii* DVIT2027 with their corresponding regions in three additional accessions: 0957 and BL-1, for which haplotype-resolved genomes are available^50^, and BS-40, represented by a haploid genome assembly^51^. We extracted the genomic regions spanning *Ren6* (PN9-057 to VMC4h4.1) and *Ren7* (VMC9a2.1 to VVIu09) from the five haplotypes of accessions 0957, BL-1, and BS-40. The length of the *Ren6*-like haplotypes ranged from 903.6 to 1,245.6 kbp, and *Ren7*-like haplotypes from 681.0 to 881.6 kbp (**Figure 6c-d**; **Table S20-S21**), indicating structural and sequence variation across accessions.

**Figure 6:**
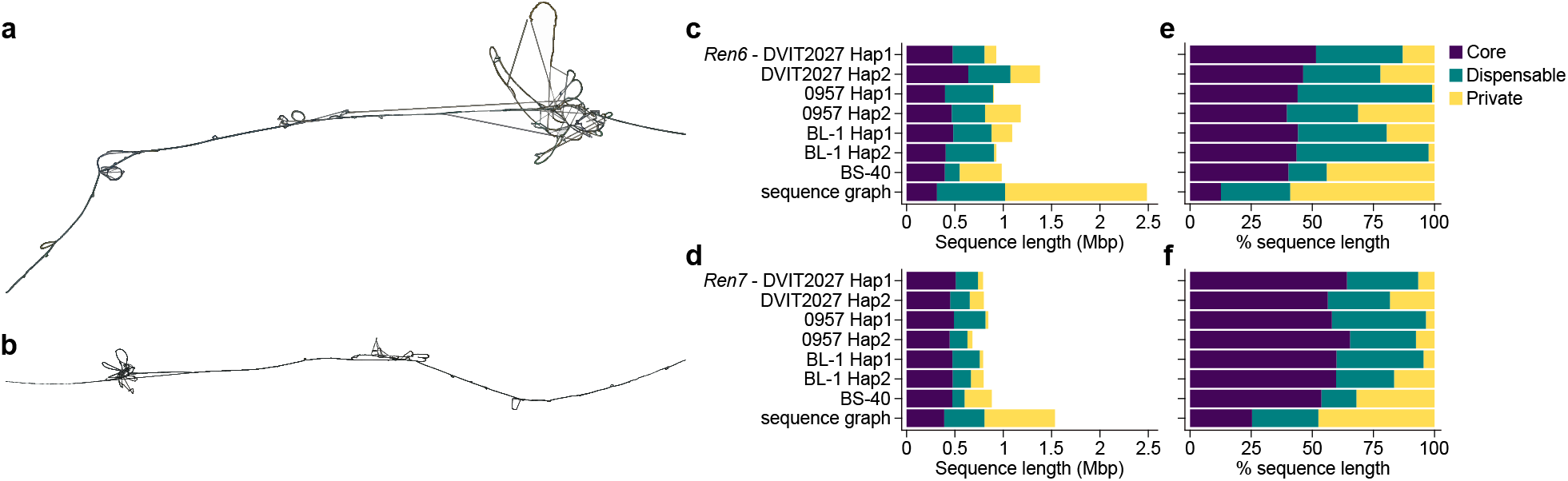
Sequence graphs of *Ren6, Ren7*, and their alternative haplotypes across four *V. piasezkii* accessions. 2D visualization of the sequence graphs built from *Ren6* (**a**), *Ren7* (**b**), and their respective alternative haplotypes in *V. piasezkii* DVIT2027, 0957, BL-1, and BS-40. Cumulative sequence length and percentage of sequence length made of core, dispensable, and private nodes in each *V. piasezkii* haplotype and the sequence graph of the genomic regions corresponding to *Ren6* (**c**,**e**) and *Ren7* (**d**,**f**). Panels **c-f** share the same legend.

To characterize this variation in greater detail, we constructed a sequence graph for each locus. The *Ren6* graph, including *Ren6* and six alternative haplotypes, comprised 107,811 nodes spanning 2,486,338 bp, while the *Ren7* graph totaled 78,411 nodes over 1,535,147 bp. These graphs capture both shared and haplotype-specific sequences and illustrate structural differences as divergent paths or “bubbles.” Prominent bubbles were observed near the end of the *Ren6* graph and the start of the *Ren7* graph (**Figure 6a–b**), consistent with large insertions and duplications in these regions.

To quantify this variation, nodes were classified into three categories: (i) core nodes, shared across all seven haplotypes; (ii) dispensable nodes, present in more than one but not all haplotypes; and (iii) private nodes, unique to a single haplotype. In the *Ren6* graph, private nodes accounted for 59% of the total graph length, dispensable nodes for 28.3%, and core nodes for only 12.7% (**Figure 6e**). Similar trends were observed in the *Ren7* graph, though the proportion of private nodes was slightly lower (47.5%) and core nodes higher (25.4%) (**Figure 6f**). These results indicate substantial sequence divergence among *V. piasezkii* accessions at both loci. The cumulative length of private nodes varied widely across haplotypes. In the *Ren6* graph, it ranged from 9.0 kbp in 0957 haplotype 1 to 433.3 kbp in the BS-40 haplotype (**Figure 6c**). A similar pattern was observed for *Ren7*, with private node lengths ranging from 29.9 kbp in 0957 haplotype 1 to 281.8 kbp in BS-40 (**Figure 6d**).

Regarding the candidate NLR genes of DVIT2027, we identified private CDS variants specific to the *Ren6* haplotype in its four candidate *R* genes: 2 bp in g158030, 27 bp in g158080, 7 bp in g158100, and 1 bp in g158430. For the *Ren7* candidates, 21 and 35 bp were found to be specific to the *R* locus in g332320 and g332350, respectively. Comparative analysis further revealed that only one NLR gene within *Ren6* (g158460) encoded a protein identical to that found in haplotype 1 of the *V. piasezkii* 0957 genome, underscoring the unique NLR composition of *Ren6* and *Ren7* in DVIT2027.

## Discussion

To better understand the genetic basis of PM resistance in wild grape species, we focused on resolving the structure, sequence diversity, and candidate genes of *Ren6* and *Ren7* in *V. piasezkii* DVIT2027. Because both loci are heterozygous in DVIT2027, it was necessary to distinguish each *R* locus from its alternative, PMS haplotype. *In silico* amplification of *Ren6* and *Ren7* associated markers revealed length polymorphisms (**Tables S20-S21**) that helped differentiate the haplotypes, although the observed lengths differed slightly from the ones previously reported^13^. However, marker-based haplotype identification alone cannot confirm locus structure or phasing accuracy. To address this, we integrated genomic data from F1 sib-lines carrying different combinations of *Ren6* and *Ren7* enabling us to verify haplotype phasing and locus structure. This required aligning sequencing reads onto both parental genomes, prompting us to also generate the diploid genome of the PM-susceptible parent of the 11373 population, F2-35. Further support to the structure of the two loci came from aligning parental-phased assemblies of six progeny back to the parental haplotypes. While *Ren6* and *Ren7*, and their PMS counterparts were fully resolved within single contigs in *V. vinifera* F2-35 and in at least one sib-line, the PMS haplotype of *Ren6* in DVIT2027 remained fragmented in all cases (**Figure 1**). These results highlight the value of phased diploid assemblies and integrated pipelines that combine long-read sequencing and high-quality phasing, while also underscoring the continued need for manual validation and careful curation of the locus structure and gene models^10^. The fragmentation observed in CLR-based assemblies reinforces the shift toward HiFi technology or other high-fidelity technologies for resolving complex *R* loci. Although the alignment of CLR reads and parental-phased genomes of the six 11373 sib-lines onto *Ren6, Ren7*, and their PMS haplotypes did not show any evidence of phasing errors or misassignments affecting the NLR gene content in the regions, undetected recombination or local assembly artifacts remain possible.

Alignments of the PMS haplotypes onto *Ren6* and *Ren7* showed large and complex SVs in the regions containing the NLR genes, as well as several short polymorphisms (**Figure 2**), similar to observations at *Run1*.*2* and *Run2*.*2* of *M. rotundifolia* Trayshed^12^. This supports a model in which NLR clusters in grapevine, as in many angiosperms, diversify at both inter- and intraspecific levels in response to pathogen pressure^62^. Phylogenetic analyses (**Figure 3**) confirmed that SVs and small polymorphisms lead to presence/absence variation, including NLRs from *Ren6* and *Ren7* without allelic counterparts in their PMS haplotypes. All NLR protein sequences in these loci were unique to the PMR haplotypes. The uniqueness of the NLR protein sequences of both *R* loci showed that sequence specificity, especially in the NLR domains, is more crucial in disease resistance rather than the number of NLRs; particularly in the case of *Ren6*, for which the PMS haplotype contains more NLRs than the PMR haplotype. This also reinforces the importance of validating phasing and structure before inferring candidate genes and highlighs the limitations of relying on reference genomes from related species or accessions that do not carry the same *R* loci^11^.

This point is further supported by our analysis of intraspecific sequence diversity at *Ren6* and *Ren7* using sequence graphs built from four *V. piasezkii* accessions. These graphs revealed substantial haplotype variation, including private sequence nodes within candidate NLRs, reinforcing the importance of generating diploid genomes from the actual resistance source. The low proportion of core sequence (12.7% in *Ren6* graph and 25.4% in *Ren7* graph) and high fraction of sequence composed of private nodes (up to 59%) indicate that most of the sequence content at these loci is accession- or haplotype-specific. Such patterns point to ongoing structural remodeling within *V. piasezkii R* loci, likely driven by pathogen selection. Notably, the BS-40 haplotype carried the largest cumulative length of private nodes, highlighting how distinct *R* locus architectures can emerge even within a single species. This divergence extended to the candidate NLR genes: several coding sequence variants were unique to the DVIT2027 haplotypes, and only one gene (g158460) had an identical protein sequence in another accession (0957). These findings underscore the unique genetic makeup of *Ren6* and *Ren7* in DVIT2027 and caution against relying on reference genomes from non-resistant accessions. Additional analyses of the accessions included in this study, and other PMR *V. piasezkii* accessions such as DVIT2026, DVIT2028, and DVIT2032^13^, could help determine whether they share PMR haplotypes and uncover additional resistance alleles or genes.

The structural variation and gene presence/absence patterns at *Ren6* and *Ren7* are likely driven by localized duplications, deletions, and transposable element (TE) activity. These mechanisms, along with unequal crossing-over, are well-known contributors to the dynamic architecture of NLR clusters^62,63^. Similar patterns have been reported at *Run1*^9^, *Run1*.*2* and *Run2*.*2*^12,20^, *Rpv3*^64^, and *Rpv33*^65^, suggesting that dynamic NLR evolution is a shared feature of grape *R* loci. With the increasing availability of phased diploid genomes from diverse *Vitis* species^10^, we are now positioned to conduct systematic comparisons across the genus. These comparisons will clarify NLR gene birth and loss, the role of TEs and recombination, and the evolutionary pressures shaping resistance loci^62,63,66^.

In addition to use genomes of 11373 sib-lines in our pipeline of identification of candidate NLR genes, we integrated transcriptomic data from leaves of six 11373 sib-lines carrying *Ren6, Ren7*, or both *R* loci, in presence or absence of the pathogen. Based on constitutive expression (**Figure 4**), and absence or sequence divergence relative to the PMS haplotype (**Figure 5**), we identified four (g158030; g158080; g158100; g158430) and two (g332380 and g332390) candidate CC-NBS-LRR genes associated with *Ren6* and *Ren7* resistance, respectively. These candidates varied in CC domain number (one or two) and number of LRR motifs (15–17), which may influence effector recognition specificity^67^. Additional transcriptomic data from recombinant lines and baseline (0 dpi) samples will help confirm and refine these candidates, especially in *Ren6*, where some genes are separated by > 500 kbp.

Future work will focus on functional validation of these candidates using transient or stable transformation methods. Transient tools available in grapevine include *Agrobacterium*-mediated expression^68,69^, virus-induced gene silencing^70^, and exogenous dsRNA-based RNAi^71^. Stable methods such as gene editing, complementation, or targeted knock-ins remain limited but are advancing^72,73^. Validating candidate NLRs and testing their response to a broader panel of *E. necator* isolates will enable development of perfect markers for breeding and guide editing strategies to stack or fine-tune resistance traits while preserving varietal identity^74^.

From a breeding perspective, the sequence-level resolution of *Ren6* and *Ren7* and their candidate genes provides a foundation for precise marker development. Given the complete haplotype specificity of these NLRs, such markers could enable rapid and accurate screening in breeding programs or guide targeted gene editing efforts^72^. The future developed markers will need to be tested in independent populations segregating for *Ren6* and *Ren7* to validate their specificity and association with PM resistance. Stacking *Ren6* and *Ren7* with other *R* loci, such as *Run1* and *Ren4*, is already underway in multiple programs to enhance resistance durability and mitigate the risk of resistance breakdown.

## Acknowledgments

This work was partially funded by the USDA NIFA Award # 2022-51181-38240 and the Ray Rossi Endowment.

## Author contributions

M.M., M.A.W., and D.C. designed the project. D.C. supervised the project. M.M., S.R., and D.P. collected the plant material. R.F.-B. extracted DNA and RNA, and prepared sequencing libraries. M.M. and N.C. performed the data analyses. M.M. and D.C. wrote the manuscript.

## Data availability

Sequencing data are accessible through NCBI under the BioProject PRJNA1258375. Genome sequences and their annotation files are available at Zenodo doi:10.5281/zenodo.15361017.

